# A nanoconfined liquid‑in‑ice aqueous regime for time-stretched single-molecule dynamics

**DOI:** 10.64898/2026.08.04.742915

**Authors:** Shao-Chuang Liu, Jia Wang, Yue-Lin Xie, Hao Chen, Ya-Xue Li, Yi-Lun Ying, Yi-Tao Long

## Abstract

The experimentally observable dynamical landscape of biomolecules is fundamentally shaped by rapid thermal motions of the surrounding aqueous environment. Although lowering temperature could expand this observable landscape, the liquid-solid phase transition of water has long prevented real-time single-molecule measurements into deeply subzero aqueous environments. Here we show that nanoconfinement within a solid-state nanopore overcomes the fundamental limitation imposed by bulk water freezing, spontaneously stabilizing a persistent liquid-in-ice environment that remains electrically accessible despite surrounding electrolyte crystallization. This aqueous environment creates a time-stretched dynamical regime, extending molecular translocation timescales by up to ∼400-fold and revealing previously inaccessible single-molecule dynamics. These findings establish a new low-temperature aqueous regime for real-time single-molecule measurements, opening new opportunities to investigate biomolecular dynamics across previously inaccessible timescales and extreme aqueous environments.

---

The experimentally accessible dynamical landscape of biomolecules is fundamentally shaped by the physical state of the surrounding aqueous environment (*1–5*). In conventional liquid water, thermal motions of water molecules continuously drive rapid molecular diffusion and frequent barrier-crossing events across rugged free-energy landscapes (*5–9*), causing transient biomolecular states to exist over exceedingly short timescales (*10, 11*). As a result, many transient molecular processes remain beyond the temporal resolution of current single-molecule techniques(*12–17*), limiting the experimentally accessible dynamical landscapes.

Lowering temperature is the most direct strategy for suppressing thermal fluctuations and extending the timescales of transient biomolecular processes into experimentally accessible regimes (*5, 10*). In cryogenic techniques such as cryo-electron microscopy, molecular motion is eliminated by freezing or vitrifying the solvent to obtain structural snapshots (*18–20*). In contrast, real-time observation of biomolecular dynamics instead requires molecules to remain continuously mobile within a liquid aqueous environment throughout the measurement. However, the liquid-solid phase transition of water has long posed a fundamental barrier to accessing deeply subzero liquid environments required for real-time studies of biomolecular dynamics. Bulk supercooled water is thermodynamically metastable and inevitably undergoes stochastic ice nucleation (*21–25*), while freezing of electrolyte solutions produces freeze-concentrated liquid regions and brine channels with heterogeneous, poorly controlled morphology and connectivity (*26–29*). Although these systems can contain residual liquid phases, they lack the long-term stability and reproducible experimental accessibility that required for real-time observation of biomolecular dynamics. Consequently, a stable low-temperature liquid aqueous regime capable of sustaining real-time single-molecule measurements has remained unavailable.

Here, we discover a nanoconfined liquid-in-ice environment within a solid-state nanopore, where a persistent electrically accessible liquid pathway is maintained within a frozen electrolyte under deeply subzero conditions, enabling real-time single-molecule measurements in a previously inaccessible aqueous state (Fig. 1). This nanoconfined liquid-in-ice environment establishes a distinct time-stretched dynamical regime in which rapid molecular processes are sufficiently slowed to bring previously inaccessible dynamic states and transitions into the observable timescale of single-molecule measurement. Within this regime, DNA translocation dynamics are extended by approximately 400-fold, shifting single-base transport from the microsecond to the millisecond timescale. This temporal expansion enables direct resolution of molecular species and transient molecular processes that remain hidden under conventional aqueous conditions. These findings open a new low-temperature aqueous regime that expands the experimentally accessible dynamical landscape of single molecules.

**Fig. 1.**
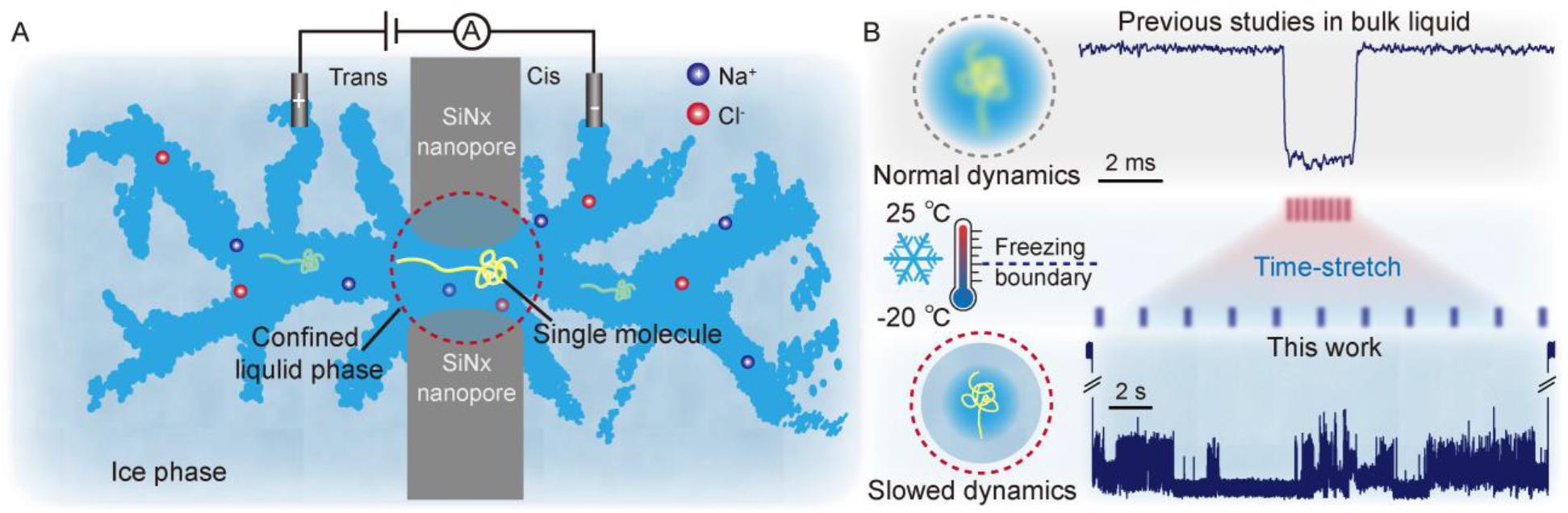
Accessing a nanoconfined liquid-in-ice aqueous regime for accessing time-stretched single-molecule dynamics. Conventional real-time single-molecule measurements are generally restricted to the bulk liquid region of the aqueous solution phase diagram, where rapid thermal fluctuations drive fast molecular dynamics. **(A)** Here, solid-state nanopore controlled nanoconfinement creates a persistent, electrically accessible liquid pathway when the surrounding electrolyte enters the frozen state, **(B)** thereby time-stretching fast molecular event into structured information-rich trajectories.

## Emergence of a nanoconfined liquid-in-ice environment under bulk electrolyte freezing

During systematic efforts to explore the low-temperature capability of nanopore single-molecule measurements, we progressively cooled a silicon nitride (SiNx) solid-state nanopore system (fig. S1) from room temperature into the subzero range (Fig. 2A). At temperature setpoints between −5 °C and −15 °C, the bulk solution underwent stochastic crystallization after variable waiting times. Unexpectedly, bulk freezing did not terminate ionic conduction through the nanopore. Instead, each crystallization event was accompanied by an abrupt increase in open-pore conductance (*G*_*0*_), while ionic current remained continuously measurable throughout the transition (Fig. 2B). The two sudden conductance increases observed during freezing are consistent with ion exclusion from the growing ice phase, resulting in solute enrichment within the remaining liquid pathway.

**Fig. 2.**
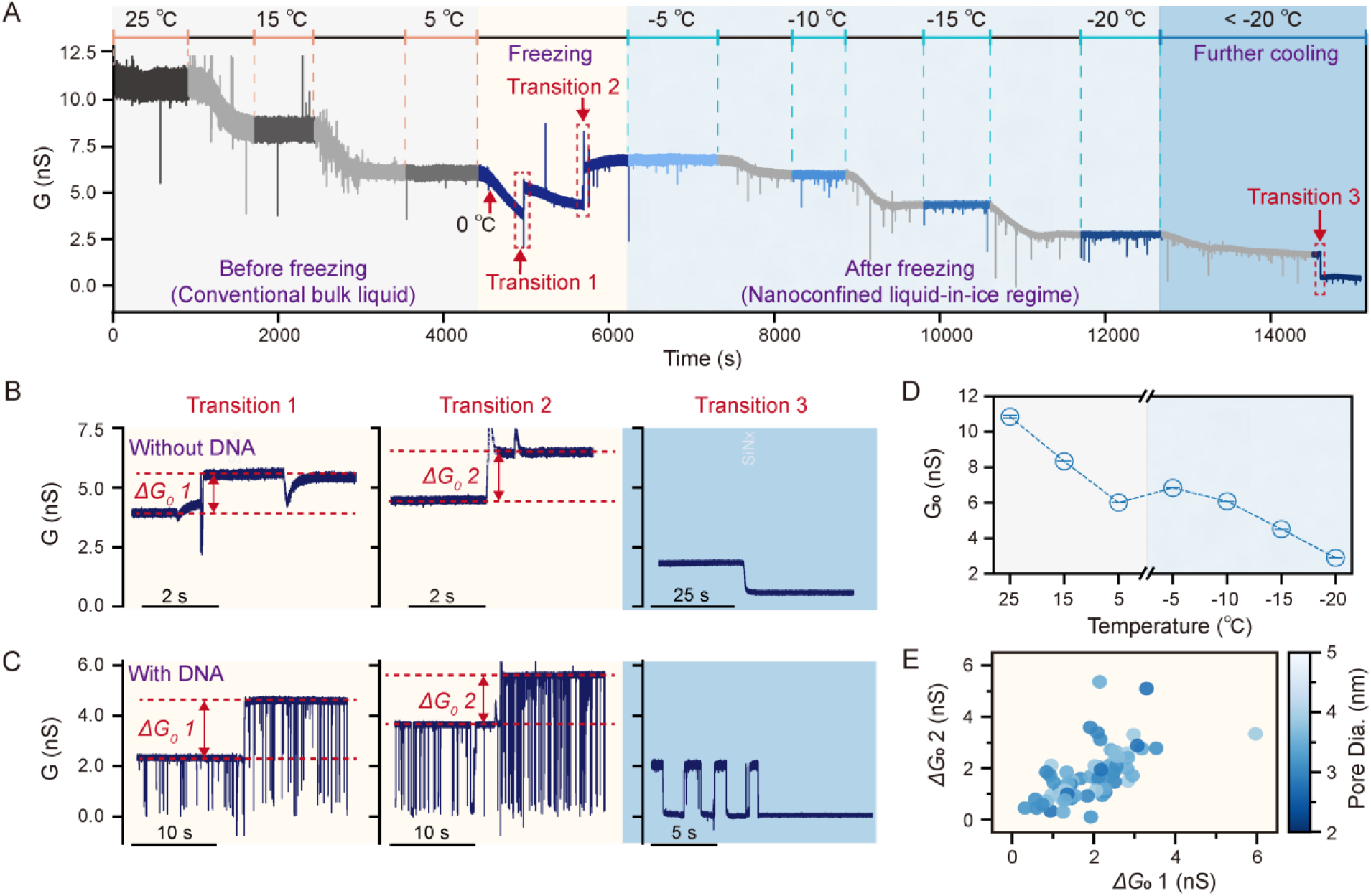
Formation of a stable nanoconfined liquid-in-ice environment in a solid-state nanopore. **(A)** Evolution of the open-pore conductance (G_0_) during controlled cooling from 25 °C to complete freezing, with the system sequentially equilibrated at fixed temperature setpoints. **(B)** The raw conductance traces of the three transition events without DNA added, showing abrupt conductance increases (ΔG_0_) associated with bulk freezing of the cis and trans reservoirs. **(C)** The raw ionic current traces of the three transition events with continuous DNA transport. **(D)** Temperature dependence of the open-pore conductance (*G*_*0*_) measured at fixed temperature setpoints during cooling. **(E)** Statistical analysis of conductance increases for the first (*ΔG*_*0*_ *1*) and second (*ΔG*_*0*_ *2*) bulk freezing transitions across nanopores with different diameters (N=72). All electrical recordings were acquired under an applied bias of +200 mV.

Following bulk crystallization, the system entered a stable and persistent conductive state that remained accessible over a broad temperature range from −5 °C to −20 °C (Fig. 2A and Fig. 2D). This conductive state remained stable over 24 hours without observable degradation, and the ionic current noise decreased systematically with decreasing temperature, providing favorable electrical conditions for low-temperature single-molecule measurements. Importantly, single-molecule DNA translocation events remained clearly detectable under these conditions (Fig. 2C), demonstrating the existence of a continuous electrically accessible liquid transport pathway within and immediately adjacent to the nanopore, despite surrounding electrolyte frozen. This liquid transport pathway exhibited remarkable stability and experimental reproducibility. Across independent nanopore (Fig. 2E) and multiple electrolyte systems, including NaCl, LiCl, and ZnCl_2_, the same transition behavior were consistently observed. The reproducibility of this liquid transport pathway contrasts with conventional freeze-concentrated electrolyte phases, where residual liquid domains emerge from heterogeneous ice growth and exhibit variable morphology and connectivity rather than a well-defined and experimentally accessible liquid pathway. These observations suggest that nanoconfinement provides a geometrically defined constraint that stabilizes an electrically connected liquid pathway within the frozen electrolyte matrix, enabling continuous single-molecule translocation through the nanopore.

We next investigated the lower temperature accessibility limit of the nanoconfined liquid-in-ice environment. As the temperature decreased below −20 °C toward the NaCl–H_2_O eutectic boundary, the persistence of the conductive state became increasingly heterogeneous across measurements. Below the nominal eutectic temperature, bulk NaCl-H_2_O phase behavior predicts the loss of a stable liquid phase and correspondingly the disappearance of continuous ionic transport. Notably, in independent measurements, the conductive pathway remained continuously accessible after cooling to −25 °C, supporting sustained ionic conduction and DNA translocation over extended measurement periods. Upon further cooling toward −30 °C, molecular transport became progressively impaired, requiring increased driving voltages and eventually becoming undetectable after prolonged operation. These observations indicate that nanoconfinement can preserve electrically addressable aqueous pathways beyond the conventional bulk eutectic temperature, while revealing a finite low-temperature accessibility limit.

Together, these results establish a robust and reproducible nanoconfined liquid-in-ice environment from −5 °C to −20 °C that maintains continuous ionic conduction and single-molecule transport within the nanopore. By providing a persistent, electrically accessible liquid pathway under bulk freezing conditions, this environment creates a new low-temperature aqueous regime for real-time single-molecule measurements beyond conventional liquid water above freezing boundary.

### Time-stretched single-molecule transport in the nanoconfined liquid-in-ice environment

We next used double-strand DNA (dsDNA) translocation as a model process to characterize the dynamical properties of this nanoconfined liquid-in-ice environment. Figure 3A show representative ionic current traces of 1 kbp dsDNA. Above the freezing transition, DNA translocation remained ultrafast and exhibited only modest temperature dependence, consistent with previous nanopore studies (*30–32*). Upon entering the liquid-in-ice environment, however, molecular transport dynamics changed abruptly. Individual translocation trajectories became progressively prolonged with decreasing temperature (Fig. 3A).

**Fig. 3.**
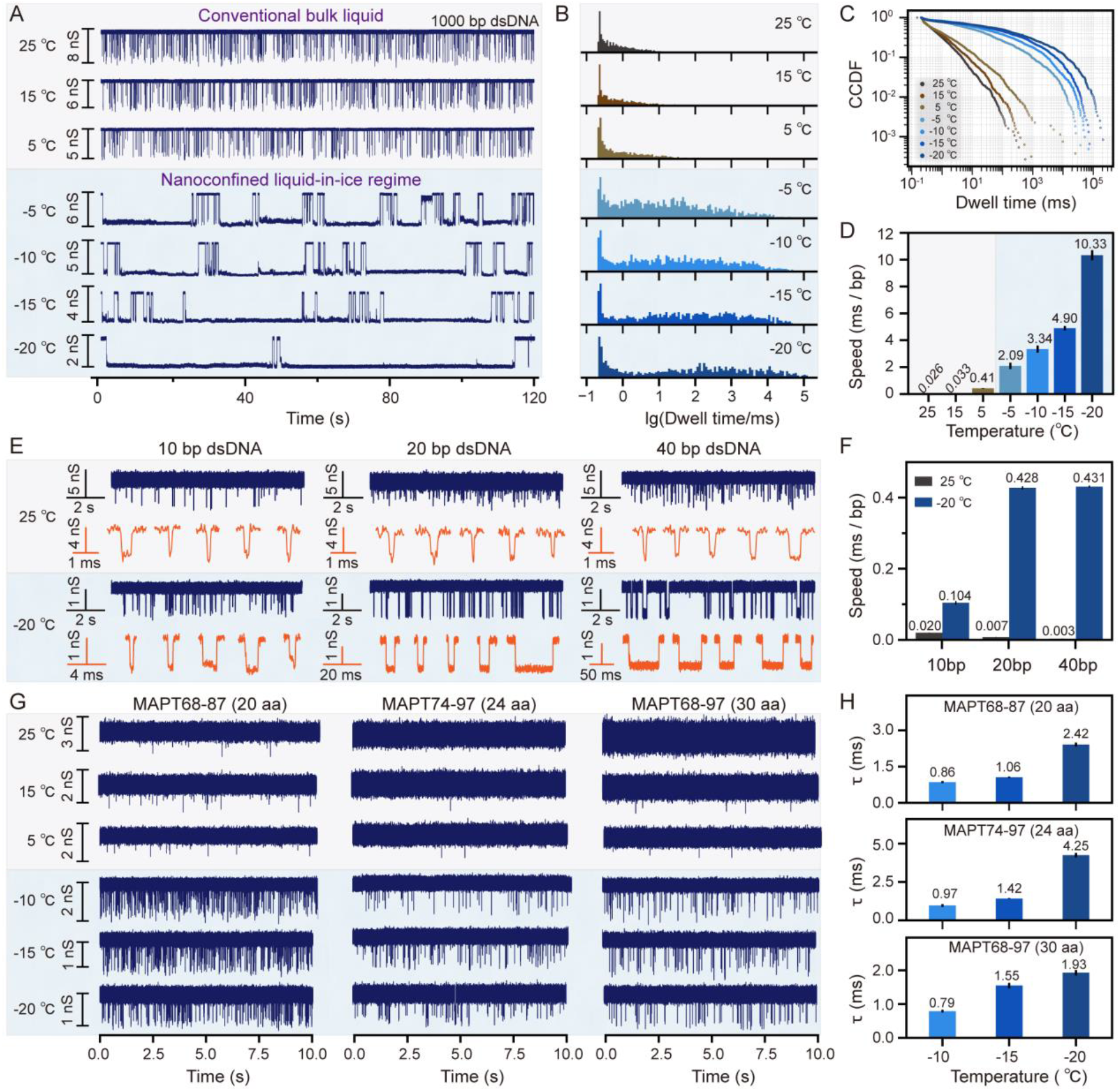
Time-stretched single-molecule transport in nanoconfined liquid-in-ice environment. **(A)** Representative ionic current traces of 1 kbp dsDNA measured before freezing and after bulk freezing at different temperatures under an applied bias of +200 mV. **(B)** The dwell-time distribution and **(C)** complementary cumulative distribution function (CCDF) plots of event dwell time for 1 kbp dsDNA translocation events (N=3719, 3176, 2647, 1637, 2045, 1517, 1193) measured at different temperature setpoints. **(D)** Temperature dependence of the characteristic dwell time per base pair (ms/bp) for 1 kbp dsDNA at + 200 mV. **(E)** Representative ionic current traces of short DNA fragments (10, 20, and 40 bp) measured at 25 °C and −20 °C under an applied bias of +100 mV. **(F)** Characteristic dwell time per base pair (ms/bp) of 10, 20, and 40 bp dsDNA measured at 25 °C and −20 °C. **(G)** Representative ionic current traces of peptide measured before freezing and after bulk freezing under an applied bias of +100 mV. Peptides include MAPT68–87, MAPT74–97, and MAPT68–97. **(H)** Characteristic translocation dwell times (τ) of peptide measured at −10 °C, −15 °C, and −20 °C .

Statistical analysis revealed that dwell-time distributions broadened substantially after freezing (Fig. 3B), while long-lived transport trajectories emerged as a pronounced long-time tail (Fig. 3C). In some cases, individual translocation events persisted for more than 100 s (fig. S33), far exceeding the timescales typically observed under ambient aqueous conditions. The characteristic translocation time per base pair (bp) increased from ∼0.026 ms/bp at 25 °C to ∼10.33 ms /bp at −20 °C (Fig. 3D), corresponding to an approximately 400-fold extension that shifted base-scale translocation from the microsecond timescale into the millisecond timescale. Notably, many time-stretched trajectories exhibited discrete current sublevels, suggesting that transient molecular conformations and molecule-pore interactions become progressively resolvable within the time-stretched dynamical regime.

Furthermore, this temporal expansion enables biomolecules that are rarely detectable under conventional aqueous conditions to become directly measurable. Short dsDNA fragments of 10, 20, and 40 bp, which normally translocate too rapidly to be reliably detected, produced well-resolved and highly reproducible blockade signals in the nanoconfined liquid-in-ice environment (Fig. 3E). At −20 °C, characteristic translocation time increased by a factor of 5, 61, and 143 for 10, 20, and 40 bp DNA, respectively, compared to measurements at 25 °C (Fig. 3F). This result indicates that time-stretching effect is length dependent for DNA, exhibiting progressively prolonged translocation trajectories for longer DNA.

To examine the generality of this time-stretching behavior across different classes of biomolecules, we next examined three small peptides comprising 20, 24, and 30 amino acids (aa). Under ambient aqueous conditions, these peptides were rarely detected because of their small excluded volume and rapid translocation dynamics (Fig. 3G). In contrast, all three peptides generated clear and reproducible translocation events within the liquid-in-ice environment (Fig. 3G). At -20 °C, characteristic dwell times (τ) reached the millisecond scale, with τ values of 2.42, 4.25 and 1.93 ms at +100 mV, respectively. These results further demonstrate that time stretching is a generalizable feature of the nanoconfined liquid-in-ice environment, enabling biomolecules that are rarely detectable under conventional aqueous conditions to be directly resolved.

Collectively, these results establish a general time-stretched dynamical regime enabled by the nanoconfined liquid-in-ice aqueous environment across diverse biomolecules. Within this regime, molecular processes that occur beyond the temporal resolution of conventional aqueous measurements are transformed into experimentally resolvable single-molecule trajectories, expanding the accessible timescale of real-time single-molecule measurements.

### Time-stretched single-molecule trajectories reveal previously inaccessible dynamics

Having established a time-stretched dynamical regime, we next asked whether extending the molecular dwell time could reveal dynamic processes that remain inaccessible under conventional aqueous conditions. We investigated a 71-nt adenine riboswitch RNA (*33*) whose translocation in nanopore requires progressive structure rearrangements (Fig. 4A). At room temperature (25 °C), adenine-bound RNA (Fig. 4B), individual trajectories remained largely featureless, appearing as nearly uniform current blockades with only poorly unresolved fluctuations (Fig. 4B). Although structural rearrangements are expected during translocation, these transitions occur too rapidly to generate distinguishable features under conventional aqueous condition.

**Fig. 4.**
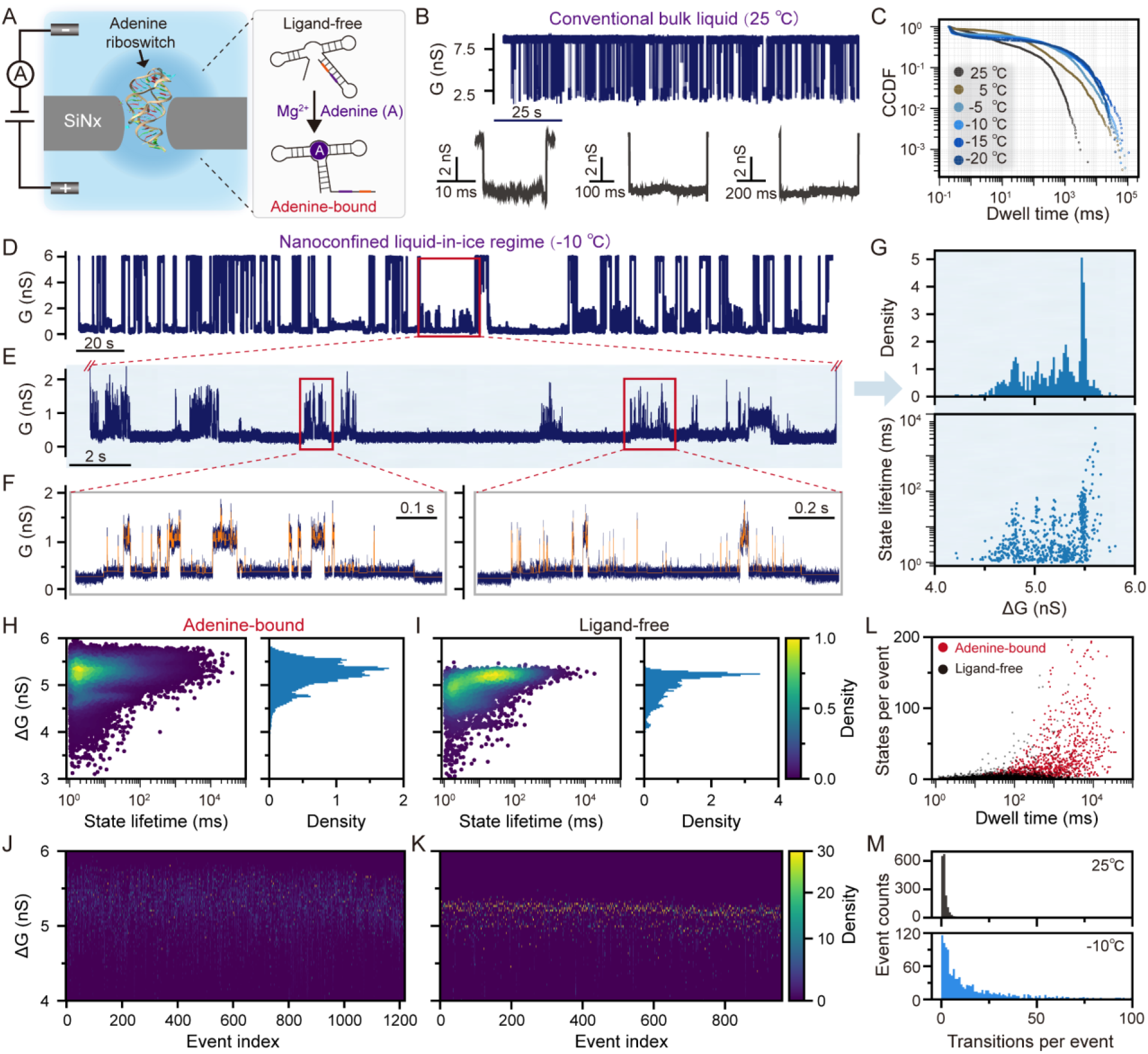
Emergence of temporally structured single-molecule trajectories. **(A)** Schematic of the experimental system showing translocation of an adenine-binding RNA (PDB code: 1Y26) through a solid-state nanopore. **(B)** Representative ionic current traces and single-molecule events of adenine-bound RNA recorded at 25 °C. **(C)** CCDF plots of event dwell times at different temperature setpoints, showing systematic time stretching upon cooling and bulk freezing. **(D)** Representative ionic current traces of RNA recorded at −10 °C after the bulk electrolyte freezing. **(E)** Representative single-molecule events at −10 °C, showing distinct conductance transitions within individual translocation events. **(F)** Magnified view of the event shown in (E). The orange line shows the fitted line by multi-level step analysis. **(G)** Scatter plot of conductance blockade (*ΔG*) versus substate lifetime and the corresponding conductance blockade histograms for the representative event shown in (E). **(H)** Ensemble substate analysis of adenine-bound RNA and **(I)** ligand-free RNA showing the global distribution of conductance blockade (ΔG) versus substate lifetime extracted from all resolved conductance substates. The point color indicates the local data density estimated by Gaussian kernel density estimation (KDE). The right panel shows the corresponding marginal histogram of conductance blockade. **(J)** Event-by-event conductance occupancy map constructed from individual adenine-bound RNA (N=1225) and **(K)** ligand-free RNA (N=960) trajectories. Each row represents one single-molecule event, and color indicates the occupancy density of each conductance blockade level. **(L)** Relationship between event dwell time and the number of resolved conductance substates per single-molecule trajectory for adenine-bound and ligand-free RNA. **(M)** Comparison of the number of resolved conductance state transitions per event for adenine-bound RNA measured at 25 °C and −10 °C. All the data show here was recorded under an applied bias of + 200 mV.

Within the nanoconfined liquid-in-ice environment, RNA translocation events extend into the tens-of-seconds range (Fig. 4C), producing a striking transformation in trajectory structure (Fig. 4D). Instead of simple featureless blockades observed under conventional aqueous conditions, individual molecular trajectories exhibited multiple discrete conductance transitions within a single translocation event (Fig. 4E). Multi-level step analysis further segmented these trajectories into a series of discrete conductance substates (Fig. 4F). As the representative trajectory shown in Figure 4E, the RNA translocation signal occupied a dominant conductance level but intermittently transitioned between distinct substates with different conductance amplitudes and lifetimes (Fig. 4G). Similar multi-state trajectories were consistently observed across individual events despite substantial event-to-event variability.

To determine whether these newly observed signal features represented reproducible single-molecule process rather than stochastic current fluctuations, we analyzed all resolved substates across the entire dataset. The resulting conductance blockade (*ΔG*)-lifetime map revealed multiple preferred conductance levels with characteristic lifetimes (Fig. 4H), demonstrating reproducible state distributions within adenine-bound RNA trajectories. In contrast, ligand-free RNA exhibited a simpler conductance landscape dominated by fewer blockade populations (Fig. 4I). These results indicate that the characteristic conductance transitions observed within individual trajectories arise from the structural dynamics of folded RNA during translocation.

Beyond ensemble-level differences, event-by-event conductance occupancy maps further revealed that individual adenine-bound RNA molecules sampled distinct subsets of conductance substates (Fig. 4J), demonstrating substantial molecule-to-molecule heterogeneity during translocation. Ligand-free RNA exhibited simpler occupancy patterns with fewer recurrent substates (Fig. 4K), consistent with a simpler dynamical landscape in the absence of ligand stabilization. These observations reveal distinct molecular dynamical landscapes at a temporal resolution inaccessible to conventional nanopore measurements, where rapid transitions are averaged within fast translocation events.

We next investigated how the expanded observation window influences the molecular information content of individual trajectories. Across individual events, the number of resolved conductance substates increased systematically with event duration (Fig. 4L), indicating that longer trajectories contain more resolvable molecular states and transitions. This increase was particularly pronounced for adenine-bound RNA, which exhibited more resolved substates and state transitions than ligand-free RNA, consistent with the greater conformational complexity of its ligand-stabilized folded structure. Further comparison found that the time-stretched adenine-bound RNA trajectories recorded in the nanoconfined liquid-in-ice environment exhibited significantly more resolved conductance state transitions than those measured at room temperature (Fig. 4M). These results demonstrate that this time-stretched dynamical regime does not merely prolong molecular signals, but expands the temporal dimension in which molecular dynamics can be observed, unfolding processes that remain compressed within conventional measurement timescales.

Collectively, these results demonstrate that the time-stretched single-molecule trajectories enable molecular processes that are too rapid to resolve under conventional aqueous conditions to become experimentally accessible. Although individual conductance substates cannot yet be uniquely assigned to specific conformational intermediates at present, their reproducible emergence reveals previously inaccessible intra-event molecular dynamics encoded within individual time-stretched translocation events.

## Discussion

This work establishes a previously inaccessible low-temperature aqueous regime by revealing that nanoconfinement of a solid-state nanopore can stabilize a persistent liquid-in-ice environment beyond the conventional freezing limitation of bulk water. Unlike cryogenic approaches that preserve molecular structures by arresting molecular motion, this regime maintains a dynamically active aqueous environment that enables continuous real-time single-molecule measurements under deeply subzero conditions. Within this environment, biomolecular dynamics exhibit an intrinsic time-stretching effect, transforming ultrafast molecular processes that are inaccessible under conventional conditions into experimentally resolvable single-molecule trajectories. Rather than slowing molecules through external manipulation of pore geometry(*34*), external fields(*35, 36*), or molecular engineering(*37, 38*), the temporal expansion achieved here emerges from reshaping the aqueous environment surrounding biomolecules to intrinsically stretches the temporal evolution of biomolecular processes, providing direct access to dynamical landscapes beyond the temporal resolution of existing single-molecule techniques.

The establishment of this aqueous regime relies on the ability of nanoconfinement to stabilize a persistent and reproducible liquid pathway within a frozen electrolyte matrix. This behavior fundamentally differs from conventional freeze-concentrated solution, where residual liquid domains typically exhibit heterogeneous morphology and stochastic connectivity. Although direct visualization of the confined liquid pathway remains challenging, the persistent ionic conduction, continuous transport of chemically diverse biomolecules, reproducibility across independent freezing cycles, and long-term stability collectively support the formation of a stable confined aqueous state. Importantly, the observation of continuous DNA translocation below the nominal NaCl-H_2_O eutectic temperature indicates that nanoconfinement can modify aqueous phase behavior beyond the equilibrium limits predicted for bulk electrolyte solutions (*39–41*). Understanding the origin of this confined aqueous state will be important for defining the ultimate temperature limits of this liquid-phase and for understanding the physicochemical principles governing low-temperature confined water.

The observed time stretching in this regime likely originates from the coupled effects of low-temperature confinement and altered molecule-solvent interactions. Unlike conventional nanopore translocation in bulk aqueous solutions, molecular transport within the liquid-in-ice pathway occurs under a modified physicochemical environment (Fig. 5A-B), where electrophoretic driving (EP), and electroosmotic counterflow (EOF) and molecule-solvent interactions are simultaneously modified. Recent studies of confined water (*42–44*) suggest that nanoconfinement can reorganize hydrogen-bond networks and modify solvent dynamics. Within the framework of Kramers’ theory (*6, 45*), enhanced solvent friction and stronger coupling between molecules and their surrounding solvent provide a potential mechanism for reducing molecular transition rates and extending the intrinsic dynamical timescales. Consistent with interpretation, the additional slowing observed in D_2_O suggests that hydrogen-bond-dependent solvent dynamics contribute to the observed temporal expansion (Fig. 5C). Future characterization of confined water structure, solvent relaxation, and nanopore electrohydrodynamics will be essential for fully resolving the microscopic origin of this emergent behavior.

**Fig. 5.**
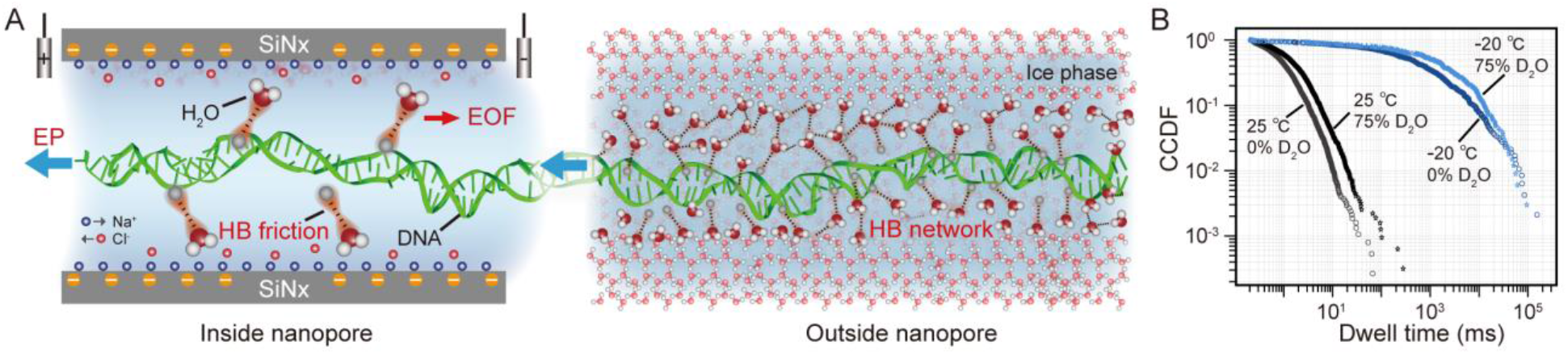
Physical framework underlying the time-stretching effect. **(A)** Schematic illustration of the electrohydrodynamic environment inside the nanopore, showing the coupled effects of electrophoretic (EP) driving and electroosmotic flow (EOF) during DNA translocation. The confined aqueous environment may enhance solvent-mediated resistance through altered water-biomolecule interactions, including hydrogen-bond-associated friction. **(B)** Schematic illustration of the surrounding frozen electrolyte matrix outside the nanopore. the confined liquid-in-ice environment may exhibit restricted water rearrangement and a more persistent hydrogen-bond network, increasing the energetic and kinetic barriers associated with molecular motion. **(C)** CCDF plots of 1 kbp DNA translocation dwell times measured in 1 M NaCl prepared in pure H_2_O and 75% D_2_O/H_2_O solvent mixtures at 25 °C and −20 °C under an applied voltage of 300 mV.

Beyond biomolecular measurements, the newly accessible aqueous regime also provides an opportunity to investigate the physical properties of confined subzero liquids themselves. In this context, individual molecules can serve as dynamic reporters of their surrounding solution environment, with molecule-solvent interactions encoded into time-stretched single-molecule trajectories. Such measurements may provide access to solvent relaxation dynamics, heterogeneity and physicochemical properties in confined aqueous phases, deeply supercooled liquids, and other nonequilibrium water environments that remain challenging to characterize directly at the single-molecule scale. More broadly, this work establishes an experimental connection between single-molecule biophysics and the study of extreme aqueous environments, opening new opportunities to explore how molecular dynamics emerge from interactions with their surrounding solvent.

## Notes

### Competing Interest Statement

The authors have declared no competing interest.

